# A multi-scale sub-voxel perfusion model to estimate diffusive capillary wall conductivity in multiple sclerosis lesions from perfusion MRI data

**DOI:** 10.1101/507103

**Authors:** Timo Koch, Bernd Flemisch, Rainer Helmig, Roland Wiest, Dominik Obrist

## Abstract

We propose a new mathematical model to infer capillary leakage coefficients from dynamic susceptibility contrast MRI data. To this end, we derive an embedded mixed-dimension flow and transport model for brain tissue perfusion on a sub-voxel scale. This model is used to obtain the contrast agent concentration distribution in a single MRI voxel during a perfusion MRI sequence. We further present a magnetic resonance signal model for the considered sequence including a model for local susceptibility effects. This allows modeling MR signal–time curves that can be compared to clinical MRI data. The proposed model can be used as a forward model in the inverse modeling problem of inferring model parameters such as the diffusive capillary wall conductivity. Acute multiple sclerosis lesions are associated with a breach in the integrity of the blood brain barrier. Applying the model to perfusion MR data of a patient with acute multiple sclerosis lesions, we conclude that diffusive capillary wall conductivity is a good indicator for characterizing activity of lesions, even if other patient-specific model parameters are not well-known.

**Author summary:** The use of advanced brain imaging techniques has supported in-vivo research targeted to the integrity of the blood-brain barrier. We propose a new type of post-processing for raw image data using contrast agent perfusion simulations on the data-poor capillary scale. Combining modern simulation techniques with the clinical image data allows us to determine patient-specific and pathologically relevant parameters such as the capillary wall conductivity. The presented simulation model is a step towards the quantification of contrast agent leakage in the brain, which is typical for acute multiple sclerosis lesions, but also occurs with other diseases affecting the blood-brain-barrier, such as cerebral gliomas.

## Introduction

Multiple sclerosis (*MS*) is characterized by a cascade of inflammatory reactions that result in the formation of acute demyelinating lesions (*MS plaques*). Acute lesions are associated with an impaired blood-brain-barrier (BBB) [1]. In healthy brain tissue, the tight junctions between endothelial cells forming the blood vessel walls, are an efficient barrier for most molecules in the brain capillaries. In active MS lesions tight junctions have been found to be damaged or open [2]. Due to an auto-immune reaction, immunological cells can pass the BBB and attack the myelin sheath covering the electrical pulse conducting axons, leading to dysfunctions of the central nervous system [3]. Magnetic resonance (*MR*) enhancement, using contrast agents such as Gadolinium-based molecules, corresponds to areas of inflammation and contrast agent leakage into the extra-vascular space. Furthermore, it is related to the histologic age of the plaques [4]. Advanced imaging techniques, such as perfusion MR imaging (*perfusion MRI*), aim at the characterization of the temporal evolution of enhancing lesion formation in relapsing-remitting MS [5]. Perfusion MRI is sensitive to inflammatory activity and can depict active lesions previous to Gadolinium enhancement and even after its disappearance [6]. Furthermore, it has been shown that perfusion in lesions is highly dynamic and related to the activity and temporal evolution of the lesions [7, 8]. Cross-sectional studies in normal appearing white matter (*NAWM*) have also demonstrated abnormal perfusion behavior in patients with MS compared with healthy controls (for review, see [9]).

Dynamic susceptibility contrast MRI (*DSC-MRI*) has proven to be informative when assessing the integrity of the blood-brain barrier (BBB) [10, 11]. In a typical DSC-MRI study, contrast agent is administered intravenously (bolus injection) and whole brain MR image sequences are recorded with a repetition time of about two seconds over a few minutes [11]. Normal appearing white matter is distinguished from inflammatory plaques by image contrast and differences in intensity–time curves. Using adequate post-processing techniques, qualitative assessment of leakage coefficients allows to identify contrast-enhancing lesions in an automated way [12]. Although today, perfusion MRI is not considered a standard procedure in the neuro-imaging workup of MS, it enables a classification of lesions according to parenchymal leakage of an MR contrast agent due to differences in perfusion behavior [13]. Perfusion imaging, both DSC and dynamic contrast enhanced (*DCE*), may provide information about the leakiness of the tissue under investigation. In this work, we investigate DSC-MRI. However, the extension of the method to DCE-MRI is conceivable.

For the interpretation of images obtained in a DSC-MRI study, the gray scale image sequence is post-processed to provide indicators within regions of interest to the radiologist. Two typical signal intensity-time curves from the brain white matter, with the characteristic first pass signal dip, are shown in Figure 1. Mathematical models (*forward model*) for contrast agent perfusion in the brain tissue can help understanding the underlying reasons for a particular intensity–time curve of a voxel, by identifying and analyzing the model parameters which are able to reproduce the MRI data. This process is also known as solving the *inverse problem*. To this end, the model parameters are tuned by using parameter estimation techniques. Forward models are typically based on a two-compartment pharmacokinetic tracer model and are parameterized by a small number of parameters [14–16]. Figure 2 visualizes a two-compartment model conceptually, with compartments representing plasma and extra-vascular, extra-cellular space. The plasma compartment is supplied by a flux determined by an arterial input function (*AIF*) [17]. The AIF can be estimated from voxels that are mostly constrained within a larger afferent artery [18]. The plasma compartment exchanges mass with the extra-vascular, extra-cellular space proportionally to its permeability-surface product. Common indicators derived from such models are the cerebral blood volume (*CBV*), the cerebral blood flow (*CBF*), the mean transit time (*MTT*), and leakage coefficients [10, 12, 19].

**Fig 1.**
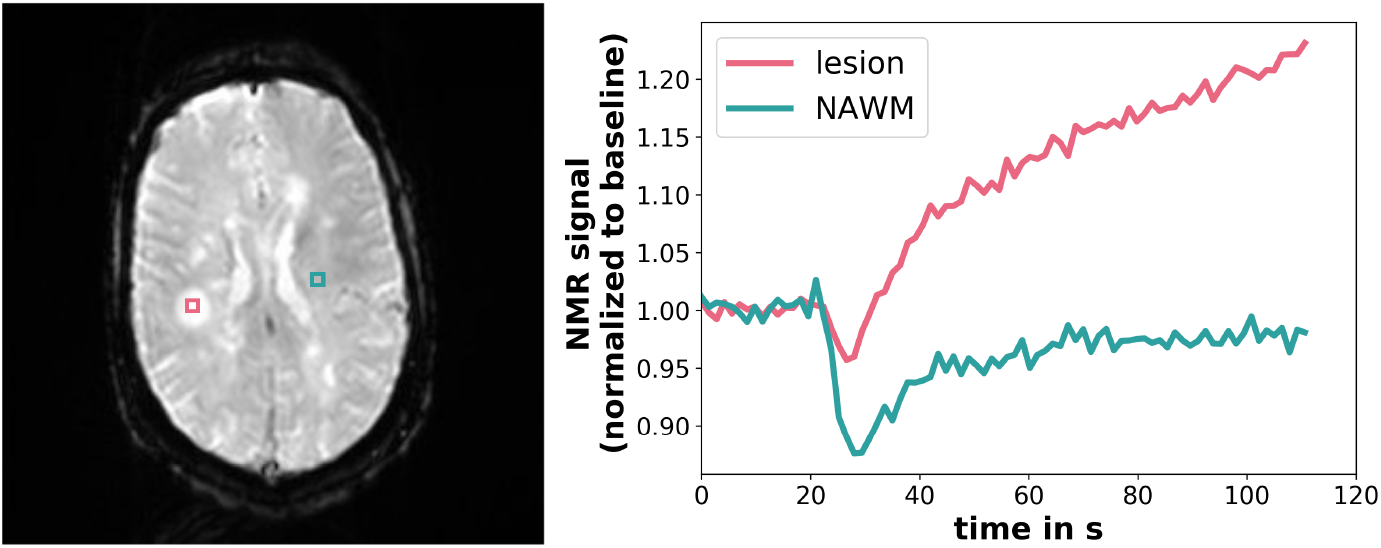
Signal intensity–time curves in a contrast-enhancing lesion (red) and in NAWM (blue) with the respective sampling locations in the brain (left). Signal values are normalized to the pre-contrast baseline. Data obtained by gradient echo - echo planar imaging (*GRE-EPI*), at magnetic field strength 3T, repetition time *T_R_* = 1400 ms, echo time *T_E_* = 29 ms, flip angle *α* = 90°, voxel size 1.8 × 1.8 × 5 mm, and an image resolution of 256 × 256 pixels per slice.

**Fig 2.**
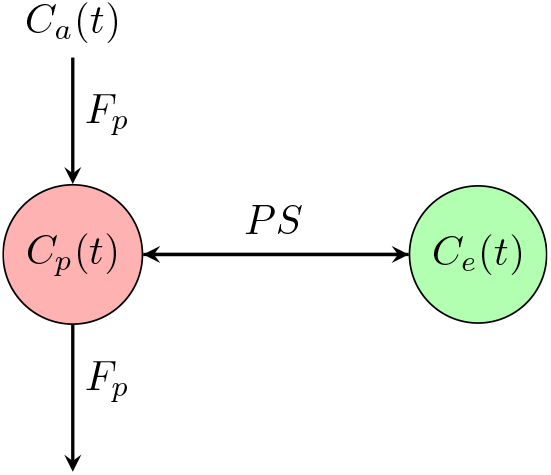
Schematic figure of a two-compartment pharmacokinetic model for tissue perfusion. Concentrations are denoted by *C*(*t*), where the subscript *a* stands for *aterial, p* for *plasma,* and e for *extra-vascular, extra-cellular space, F_p_* is the plasma flux, and *PS* denotes the permeability-surface product, a proportionality constant of the transmural exchange rate.

A routinely used state-of-the-art post-processing procedure and model is described in [12]. Such models have to reflect two processes: (1) the perfusion process governed mainly by bio-fluid-mechanical principles, and (2) the physical process of nuclear magnetic resonance (*NMR*) exploited to acquire the MR image. There have been many suggestions for improving the modeling of the latter process [20–23]. The authors of [24, 25] show that the local, sub-voxel tissue structure has a significant effect on the NMR signal. However, all previous studies, including the recent study by [23], rely on state-of-the-art two-compartment models for the perfusion process providing only average concentrations in two tissue compartments within a voxel.

To overcome the limitations of two-compartment models, we present a perfusion model on a sub-voxel scale, including the capillary network structure. Fully, three-dimensionally resolved fluid-mechanical models of brain tissue perfusion imply prohibitively complex and computationally expensive simulations due to the large number of vessels, their non-trivial geometrical embedding, and the complex geometry of the extra-vascular, extra-cellular space [26]. To reduce complexity, we use a mixed-dimension embedded model description, where blood vessels are represented by a network of cylindrical segments which are embedded into the extra-vascular space, represented by a homogenized three-dimensional continuum. The model reduction, which is described in more detail in the following, leads to a coupled system of one-dimensional partial differential equations for flow and transport in the vessels, and three-dimensional partial differential equations for flow and transport in the extra-vascular space. Related models have been used to study the proliferation of cancer drugs [27–29], the transport of oxygen [30–34], and nano-particle transport for hypothermia therapy [35]. A recent study [36] describes contrast agent perfusion based on diffusive transport with a mixed-dimension model. The herein presented fluid-mechanical model is similar to the drug proliferation model described in [27] and introduced by [28]. It is derived here for the specific application of contrast agent perfusion in brain tissue.

The fluid-mechanical model is coupled to an NMR signal model. We propose that the local distribution of the contrast agent and resulting local susceptibility effects obtained by a sub-voxel scale model may better explain the NMR signal response of the tissue. The application of this new perfusion model is demonstrated for the example of MS lesions.

In the following, we refer to the sub-voxel spatial scale, ranging from a few micrometers to several hundreds of micrometers, as *meso-scale*. We call the scale below the meso-scale, which includes the molecular scale, *micro-scale*, and refer to the scale above as *macro-scale*.

## Mixed-dimension embedded model for brain tissue perfusion

The tissue is conceptually decomposed into two domains. The *vascular* compartment comprises blood vessels, including the capillary lumen, the endothelial surface layer, the basement membrane, and blood. The *extra-vascular* compartment includes cells, the extra-cellular matrix (*ECM*), and the interstitial fluid. The compartments communicate by the exchange of fluids and molecules over the capillary wall (*transmural exchange*). In the following three sections, the assumptions are discussed separately for both compartments and the transmural exchange. These sub-domain models are then combined, to obtain the mixed-dimension tissue perfusion model.

### Vascular compartment

Blood flow can generally be described by the Navier-Stokes equations. Assuming negligible radial velocities, long vessels (compared to their radius), low Reynolds numbers (*Re* << 1), non-pulsatile flow, and rigid vessel walls, the equations can be simplified [37]. Furthermore, we employ a homogenized continuum model for blood, using an apparent viscosity, 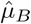. Assuming a constant hematocrit of 45%, the apparent viscosity can be described as a function of the effective vessel radius, *R*, by an empirical relation [38], derived from experimental data,

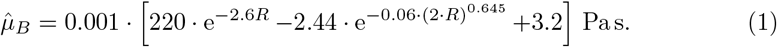

The effective vessel radius, *R*, is chosen with respect to the cross-section area, *A_v_* = *πR*^2^. Blood density is assumed constant, *ρ_B_* = 1050 kg m^-3^ [39]. Under these assumptions, the flow in the lumen of a capillary vessel can be described by

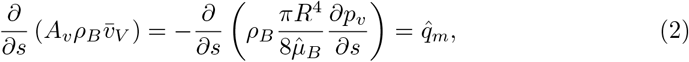

with the mean velocity 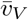, the rate of mass exchange with the extra-vascular compartment, 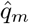, with units of kgs^-1^ m^-1^, the cross-section averaged pressure, *p_v_*, and the axial coordinate, *s*. At vessel bifurcations, we enforce continuity of pressure and conservation of mass to couple the equations of the different vessel segments.

The transport of contrast agent can be described by an advection-diffusion equation. By integration of the three-dimensional equations over the vessel cross-section, the model can reduced to a one-dimensional equation for the cross-section-averaged mole fraction, *x_v_*, [37]

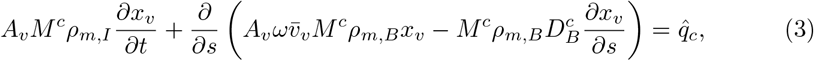

where *M^c^* is the molar mass of the contrast agent, *ρ_m,B_* the molar density of blood, and *D^c^_B_* the binary diffusion coefficient of the contrast agent in blood. The exchange with the extra-vascular compartment is modeled by the flux 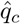, with units of kgs^-1^ m^-1^. The shape factor *ω* > 0 reflects the variation of axial velocity profiles in vessel cross-sections [37],

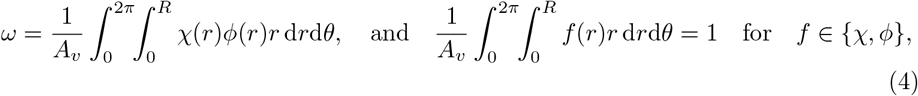

where *χ*(*r*), *ϕ*(*r*) are the dimensionless velocity profile and the dimensionless concentration profile, respectively. As it has been observed that small nano particles are likely to be distributed evenly [40], we choose *ω* = 1.

In the following, we consider the Gadolinium-based contrast agent Gadobutrol. For the perfusion MRI sequence, it is administered intravenously in solution, with a concentration of 1 moll^-1^. Gadobutrol has the chemical formula C_18_H_31_GdN_4_O_9_, corresponding to a molar mass of *M^c^* = 604.715 g mol^-1^ [41]. In high concentrations, Gadobutrol has a significant influence on fluid density and viscosity. However, the concentrations arriving in the brain tissue sample are strongly diluted by diffusion and dispersion along the tortuous path through the vascular network, so that the influence on blood density and viscosity can be neglected in this study. The binary diffusion coefficient of Gadobutrol in plasma can be estimated by means of the Stokes-Einstein radius, *r*_hy_ = 0.9nm [42],

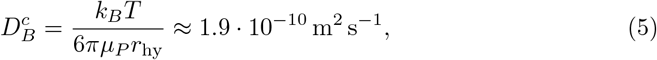

where *μ_P_* = 1.32Pa s [43] denotes the blood plasma viscosity, *T* the temperature in K, and *k_B_* the Boltzmann constant.

### Extra-vascular compartment

The extra-vascular compartment is modeled as a porous medium with a rigid solid skeleton, consisting of cells, fibers, and extra-cellular matrix. Flow of a single fluid phase, the interstitial fluid, through a porous medium is described by Darcy’s law [44]

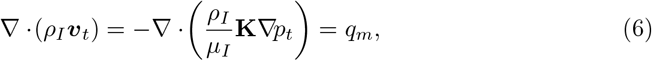

where *ρ_I_*, *μ_I_* are density and viscosity of the interstitial fluid, ***v**_t_* the filter velocity vector, **K** the intrinsic permeability tensor of the extra-vascular compartment, and *q_m_* (kg s^−1^ m^−3^) the mass exchange with the vascular compartment. We assume constant density and viscosity, *ρ_I_* = 1030 kg m^−3^, *μ_I_* = 1.32Pa s, given that contrast agent concentrations in the extra-vascular compartment are even smaller than in the blood stream, and we consider perfusion an isothermal process. Furthermore, we choose an isotropic intrinsic permeability **K** = 8.3 · 10^−18^ m^2^ [45], where **K** = *K***I**. The transport is modeled by an advection-diffusion equation,

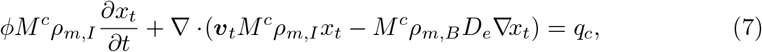

where *ϕ* denotes the porosity, the ratio of pore volume to total volume in a representative elementary volume, *ρ_m,I_* the molar density of the interstitial fluid solution, *D_e_* is the effective diffusion coefficient, and *q_c_* (kg s^−1^ m^−3^) is the contrast agent mass exchange with the vascular compartment. We assume that the interstitial space in the extracellular matrix, with pore throat diameters of around 50 nm [26], still allows for a viscous flow regime. Furthermore, it is assumed that Gadobutrol will not enter cells. The effective diffusion coefficient in the porous medium can be estimated as 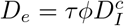, where *τ* denotes the tortuosity of the extra-cellular matrix, and 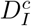 the binary diffusion coefficient of contrast agent in interstitial fluid, for which we choose the same value as for the binary diffusion coefficient of contrast agent in plasma Eq. (5). Following the literature for tortuosity and porosity values [26], we choose τ = 0.4 and *ϕ* = 0.2, which yields, *D_e_* ≈ 1.5 · 10^−11^ m^2^ s^−1^.

### Transmural exchange

The wall of continuous capillaries consists of an endothelial surface layer, a basal membrane, and a layer of charged proteins, called glycocalyx [46, 47]. Mass exchange can occur passively through the endothelial tight junctions, or through trans-cellular pathways. Here, we consider only transport by advection and diffusion, following [39]. Given a blood vessel volume fraction of 3 %, an average thickness of the endothelial surface layer of 1 μm [48], and an average vessel radius of 10 μm, the volume fraction of the capillary wall is less than 1 % of the tissue volume. The capillary wall can be conceptually reduced to a two-dimensional interface, denoted by Γ, separating the vascular from the extra-vascular compartment. Note that this results in a pressure jump across Γ, which is inversely proportional to wall permeability and wall thickness. According to Starling’s hypothesis [49, 50], the transmural flux of a fluid is proportional to the hydrostatic and colloid osmotic pressure gradient between capillary lumen and interstitial space

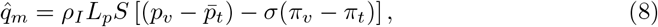

where *L_p_* is the *filtration coefficient*, with units of m Pa^−1^ s^−1^, *S* = 2*πR* is the circumference of the vessel,

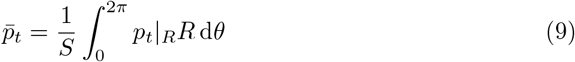

is the average hydrostatic pressure on the vessel wall, *π_v_, π_t_*, denote the osmotic pressure in capillary lumen and interstitial space, respectively, and 0 ≤ *σ* ≤ 1 is the osmotic reflection coefficient. The difference in osmotic pressure results from large plasma proteins in the blood stream, such as *albumin*, and effectively draws fluid into the vessels. For the in silico experiments, we assume the osmotic pressures to be constant, with Δ*π* = *π_v_* − *π_t_* = 2633 Pa [51]. Furthermore, we choose σ =1, corresponding to the vessel wall modeled as a perfect selectively permeable membrane.

The contrast agent is assumed to be transported by advection with the plasma, as well as by molecular diffusion. The reduction of the vessel wall to a surface leads to a concentration jump across the vessel wall, which is inversely proportional to diffusive wall conductivity and wall thickness. The transmural transport is described as [50]

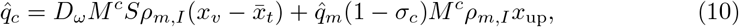

where *D_ω_* is the diffusive wall conductivity, with units of ms^−1^,

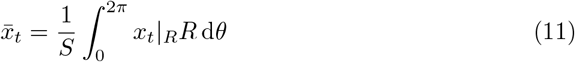

is the average contrast agent mole fraction on the vessel wall,

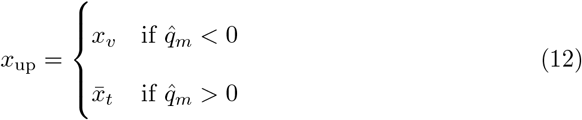

denotes the mole fraction in upwind direction, and 0 ≤ *σ_c_* ≤ 1 denotes the solvent-drag reflection coefficient. As the considered contrast agent is a small molecule and the endothelial tight junctions are damaged in lesion tissue, we set *σ_c_* = 0, neglecting reflection. Determining *D_ω_* from MRI data is the major objective of this work.

The mass balance Eqs. (2), (3), (6) and (7) are coupled by Eqs. (8) and (10), whereas Eqs. (6) and (7) are described in the three-dimensional extra-vascular domain Ω, while Eqs. (2) and (3) are associated with the one-dimensional vascular domain Λ. We follow the concept suggested in [28]: if the source terms, 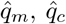, are defined as line sources along the vessel center line in the three-dimensional domain, while the three-dimensional quantities, *p_t_, x_t_*, are evaluated as the average values on Γ, then, the resulting exchange term is a good approximation of the source term in a non-reduced three-dimensional setting. To this end, we define *q_m_*, the source term in Eq. (6), as

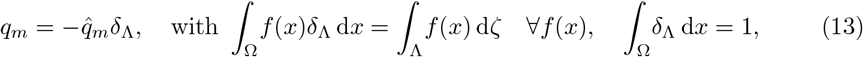

so that *q_m_* is a line source restricted by the Dirac delta function *δ*_Λ_ to the center line of a vessel. Analogously, we set 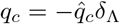, for the source term in Eq. (7).

### Vessel geometry, boundary conditions and initial conditions

We base our vascular model on a small network of capillaries from the superficial cortex of the rat [34, 52], which we consider a sufficient approximation of the actual capillary network geometry for type of model analysis presented in this work. The network has the dimensions 150 μm × 160 μm × 140 μm, and is shown in Fig. 3. The location of inflow and outflow boundaries are given in this data set. For the inflow boundaries, [34] provide velocity estimates based on the vessel radius, which are applied as Neumann boundary conditions. At the outflow boundaries, we enforce Dirichlet boundary conditions for the pressure, *p*_*v*,out_ = 1.025 · 10^5^ Pa.

**Fig 3.**
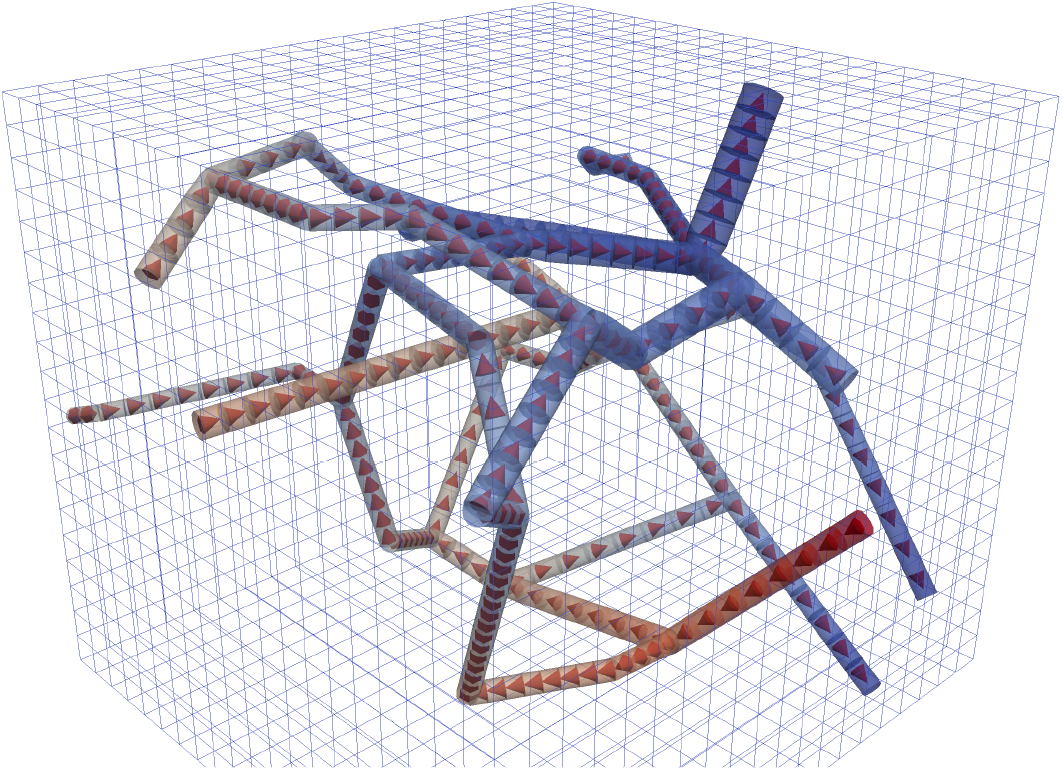
The capillary network grid extracted from measurements in the rat cortex [34, 52] and the Cartesian computational grid for the extra-vascular domain, used for the model analysis in this study. The tubes are scaled with the respective vessel radius. The color visualizes hydraulic pressure from high (red) to low (blue). The cones indicate the flow direction.

The domain initially contains no contrast agent, so that *x_v_*(*x, t*) = 0. During the perfusion MR study, 10 ml contrast agent (0.1 mmol per kg body weight) is administered intravenously as a solution at 5 mls^−1^ and a concentration of 1 moll^−1^. The injected fluid thus forms a sharp bolus. However, the bolus disperses significantly before it reaches the brain capillaries. Therefore, the concentration inflow profile to the capillary network has to be estimated from the parameters of the bolus injection. To this end, we use an ansatz from [22]

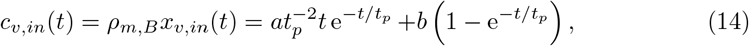

which describes a concentration profile starting at *c*_*v*,in_(0) = 0molm^−3^ and approaching an equilibrium concentration *b* (molm^−3^, contrast agent is equally distributed in the whole body blood volume), with a single peak after the arrival of the bolus. The parameters *a* (mol sm^−3^) and *t_p_* (s) are shape parameters of the capillary input function, and can be interpreted as the scaling parameter for the area under the curve, and the time to peak, respectively, in the absence of re-circulation (*b* = 0). The parameter values are patient-specific and also depend on the location in the brain. Values for *a,b*, and *t_p_* are discussed below, in the context of parameter estimation.

At the inflow boundary, contrast agent influx is enforced by a Neumann boundary condition. At the outflow boundary, the normal mole fraction gradient is set to zero and the advective component flux is computed by a first-order upwind scheme. For the extra-cellular compartment, we enforce symmetry boundary conditions everywhere, assuming that the modeled domain is surrounded by domains with similar properties.

### Mixed-dimension embedded model for tissue perfusion

In summary, the complete coupled fluid mechanical model of tissue perfusion reads as:

1. Find *p_t_,p_v_* such that

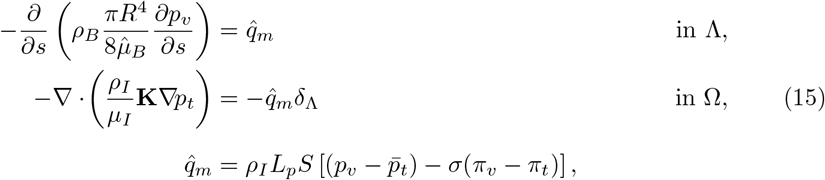
2. then find *x_t_,x_v_* such that

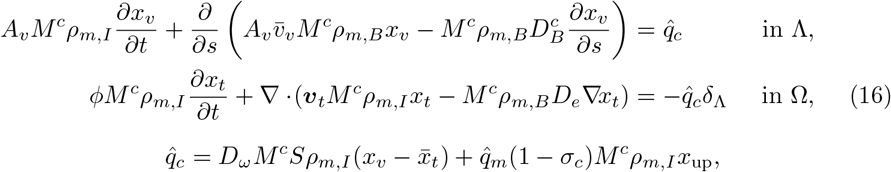

subject to the Neumann and Dirichlet boundary conditions on the inflow and outflow boundaries of the vascular compartment, *∂*Λ, respectively, and no-flow boundaries for the boundaries of the extra-vascular domain, *∂*Ω, as discussed in the previous section.

This model stands in contrast to the often employed two-compartment kinetic modeling approaches, because it resolves meso-scale flow phenomena, and because it is based on parameters with a clear physical interpretation.

## NMR signal model

A model linking concentration fields with the nuclear magnetic resonance (*NMR*) signal response is required to connect the results of the fluid mechanical model to clinical MRI data. To this end, we develop a model of NMR on the meso-scale. In the following, we describe a gradient echo, echo planar sequence (*GRE-EPI*) commonly used in DSC-MRI. This fast imaging technique allows acquisition of an entire brain image stack in less than two seconds. Thus, after the injection of a contrast agent, a time series of such images can be acquired, where the characteristic signal-time curve for every voxel is dependent on the evolution of the contrast agent concentration distribution on the meso-scale.

The GRE-EPI sequence starts with a radio frequency (*RF*) pulse, which reorients the magnetic moments in the tissue sample, with the flip angle *α* to the main magnetic field ***B*_0_**. The RF pulse causes the magnetic moments to precess. Energy dissipation, characterized by an exponential decay with the longitudinal and transversal relaxation times, *T*_1_, 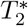, relaxes the magnetization into the initial state aligned with ***B*_0_**. According to [22], the GRE-EPI voxel signal during a DSC-MRI perfusion sequence can be modeled as

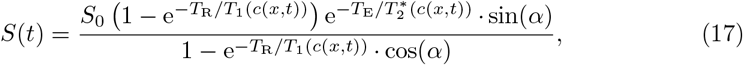

where the repetition time *T*_R_, is the time between two RF pulses, and the effective echo time, *T*_E_, is the time between RF pulse and signal readout. The base signal *S*_0_ > 0 depends, i.a., on tissue proton density and the MR scanner hardware. In the following, we look only at the normalized signal 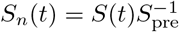, where *S*_pre_ is the signal before the contrast agent bolus arrives in the tissue sample. The pre-contrast signal, *S*_pre_, contains all constant factors in Eq. (17), including *S*_0_. It follows from Eq. (17) that a shortening of 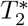 results in a decrease of NMR signal strength, while a shortening of *T*_1_ results in signal enhancement.

The following two sections introduce the models for the relaxation rates 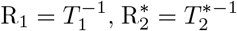 that depend on the spatial and temporal evolution of the contrast agent concentration, *c*(*x,t*), within an MRI voxel. The authors of [23] developed a model including an artificial microstructure using a combination of a finite perturbator method [21] and a finite-difference solution of the Bloch-Torrey equations. However, their model is coupled to a two-compartment tracer perfusion model, only providing voxel-averaged concentrations. In contrast, the presented perfusion model computes the sub-voxel distribution of the contrast agent concentration. We follow [22], to develop a model considering the spatial and temporal distribution of the contrast agent.

### Transversal relaxation in tissues with locally heterogeneous microstructure

The transversal relaxation rate, 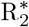, depends on the complex local microstructure of the tissue [24] and is altered by the presence of the contrast agent. We are only interested in the signal change relative to the baseline, so we split the relaxation rate in a static pre-contrast contribution and a time-dependent contribution depending on the contrast agent concentration,

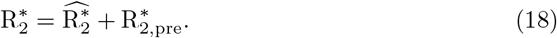

The relaxation rate for a sub-voxel control volume can be described by contributions of three compartments, the vascular compartment (B), the extra-cellular, extra-vascular space (I), and the cellular compartment (S), weighted by their volume fractions, *ϕ*_B_, *ϕ*_I_, *ϕ*_S_ [22],

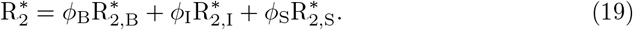

According to [25], the rate in each compartment *κ* ∈ {B, I, S}, comprises contributions on three spatial scales

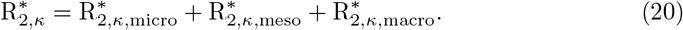

The rate 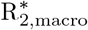 describes effects of static local inhomogeneities of the magnetic field ***B*_0_**, which are time-independent. Since the static effects do not depend on the contrast agent concentration, they are included in the pre-contrast relaxation rate, 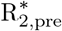. The rate 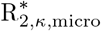 depends on the local chemical composition. The effects are independent of the pulse sequence. Gadolinium-based contrast agent molecules increase the relaxation rate, which can be described by a linear relationship [25],

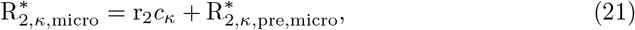

where r_2_ is the molar relaxivity, and *c_κ_* the local contrast agent concentration in compartment *κ*. The molar *T*_2_ relaxivity, r_2_, of Gadobutrol at 3T and 37°C is approximately 3.9 m^3^ mol^−1^ s^−1^ [53]. Here, we assume that the contrast agent cannot enter the cells, *c*_S_ = 0, hence 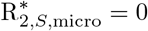.

The term 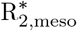 stems from a meso-scale effect. The magnetic field perturbations induced by the difference in magnetic susceptibility in the blood vessel and the extra-vascular space, increase the relaxation rate of the extra-vascular space in proximity of a blood vessel. The generated magnetic field perturbations are several orders of magnitude smaller than ***B*_0_**. Furthermore, the influence decays rapidly with distance to the vessel surface. Therefore, we consider each segment of the vessel network to cause a perturbation independent of the other segments. The increase in 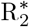 at a given location in the tissue caused by mesoscopic magnetic field perturbation will then be the superposition of all n segment perturbations

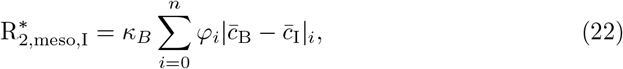

where 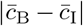 is the difference of the concentrations averaged on the vessel surface. The factor *κ_B_* ≥ 0 is an ad-hoc parameter, scaling the strength of these perturbations. The proportionality factor *φ_i_* models the decay of the influence of the with distance from the vessel wall. We set *φ_i_* = *R*^2^/*r*^2^, assuming a quadratic decay, where *r* is the distance to the vessel center line and *R* the radius of the vessel segment. The susceptibility contrast likewise increases the transversal relaxation rate, which we model by

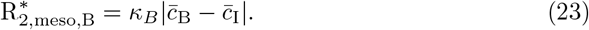

The same effect occurs at the cell surfaces, induced by the difference in magnetic susceptibility between interstitial space and cells. Note that we consider cells not to be invaded by contrast agent. We include this effect by adding a term to Eq. (22),

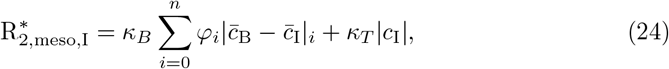

and to the relaxation rate of the cell compartment,

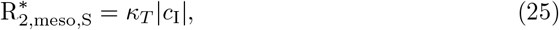

where *κ_T_* ≥ 0 is a second ad-hoc parameter, determining the strength of these perturbations. Furthermore, we assume that there is no direct interface between the cells and the vascular compartment.

Combining Eqs. (19), (21) and (23) to (25), we obtain a formulation for the transversal relaxation rate dependent on the concentration fields and the volume fractions of the three compartments:

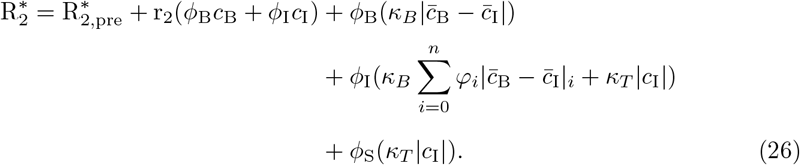

### Longitudial relaxation with contrast agent administration

Similar to 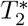, the contrast agent also shortens *T*_1_. However, the effects occur merely on the micro-scale. Thus, we can model the relaxation rate R_1_ = 1/*T*_1_ of the tissue sample by

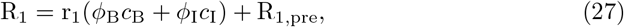

where we implicitly assumed that contrast agent does not enter cells, *c*_S_ (***x**,t*) = 0. The molar *T*_1_ relaxivity, r_1_, of Gadobutrol at 3T and 37 °C is approximately 3.2 m^3^ mol^−1^ s^−1^ [53].

### Voxel signal

The relaxation rates, 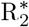 and R_1_ (Eqs. (26) and (27)), are computed for each control volume in the three-dimensional domain Ω. The volume fraction of the vascular domain, *ϕ_B_*, is computed by integrating over the volume of all vessels within a control volume and dividing this number by the volume of the control volume. A local NMR signal can then be computed for each control volume, by using Eq. (17). The voxel signal is determined by the volume average of all control volume signals.

## Numerical treatment and implementation

The equations of the fluid flow equation system (Eq. (15)), and the contrast agent transport system (Eq. (16)), are discretized with a cell-centered finite volume method with a two-point flux approximation in space, and an implicit Euler method in time. The two systems are only coupled in one direction, such that Eq. (16) depends on the pressure field computed in Eq. (15), but Eq. (15) can be solved independently of Eq. (16). Furthermore, Eq. (15) is stationary, so that the pressure field only has to be computed once per perfusion experiment. The discrete systems are assembled in a block-matrix structure in residual form,

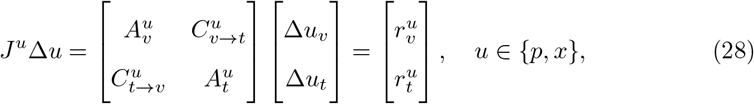

where *u* is the respective discrete primary variable (fluid pressure in Eq. (15), contrast agent mole fraction in Eq. (16)), and Δ*u* denotes the difference of the current solution to the solution of the previous time step (or initial solution). The Jacobian matrix *J^u^* can be split into blocks, where 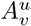 is the block with derivatives of the residual 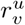 with respect to the degrees of freedom *u_v_* of the discrete vascular domain Λ_*h*_ ⊆ Λ, 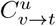 contains derivatives of the residual 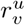 with respect to the degrees of freedom *u_t_* of the discrete extra-vascular domain Ω_h_ ⊆ Ω, and 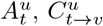 are defined analogously in terms of the residual 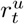.

The linear equation systems, Eq. (28), are solved using a left preconditioned stabilized bi-conjugate gradient method [54, Chapter 7], with the block diagonal perconditioner

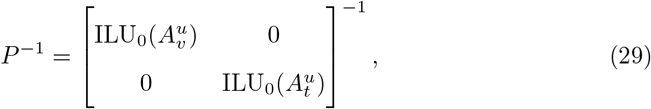

where ILU_0_(*A*) denotes an incomplete LU-factorization of the matrix *A* using *A*’s sparsity pattern (zero fill-in) [54, Chapter 10].

We assume that the influence of the sub-voxel contrast agent evolution during a single image acquisition on the NMR signal is negligible, and thus, Eq. (17) is solved as a post-processing step after each time step of the perfusion model.

The model converges in time and space to a reference solution computed on a very fine grid and a very small time step size. The convergence study is described in detail in S1 Appendix. As a result of the convergence study, we choose our computational grids such that the largest grid cell does not exceed 8 μm. This results in a run-time of a few seconds on a normal laptop for a single forward model run.

The model is implemented with the open-source porous media simulator DuMu^x^ [55], which is based on the Distributed Unified Numeric Environment (DUNE) [56, 57]. The implementation of the mixed-dimension embedded tissue perfusion model is based on a recent extension of DuMu^x^ for multi-domain porous media problems, first described in [58] for the simulation of root-soil interaction in the vadose zone. We refer to this publication for a more detailed description of the discretization, assembly procedure, and software implementation of mixed-dimension embedded models.

## Inverse modeling using clinical MRI data

We use clinical MRI data to evaluate the presented model. We choose a patient with relapsing-remitting MS from a clinical study with 12 MS patients, diagnosed according to the revised McDonald’s criteria [59], and showing at least one contrast enhancing lesion on MRI. The data is selected from a previous study that has been published elsewhere [60], and fully anonymized for further analysis. For the employed GRE-EPI protocol, 19 parallel images with a slice thickness of 5 mm are taken 80 times during an acquisition time of 119 s. The sequence parameters are given in the caption of Fig. 1. From these images, a clinical expert annotated a voxel within a Gadolinium enhancing MS lesion (sample L) and a corresponding voxel in NAWM (sample N). Fig. 1 shows the samples L and N, together with the respective voxel locations in the MRI slice.

Several model parameters can be assigned a fixed value, either because the parameter assumes a well-known fixed value given in the literature, or because the parameter is not expected to significantly affect the results of this particular study and an approximate value can be obtained from the literature. However, there are also parameters that are inherently patient-specific and cannot be directly measured, or parameters for which the measurement data is not available for the given patient. These parameters are, *a, b, t_p_, k_B_, k_T_, T*_1,pre_, 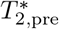, *L_p_, D_ω_*. Determining these parameters for a given signal-time curve constitutes an inverse problem. In particular, we aim to determine *D_ω_*, which may quantify contrast agent leakage, and thus, has direct clinical relevance.

In the following, we briefly discuss typical values or value ranges for these parameters. The shape parameters, *a, b, t_p_*, determine the inflow profile of the bolus arriving at the voxel under study. They are generally varying from voxel to voxel. In particular, *a* and *t_p_* depend on the voxel location and vessel network structure, as well as the resulting bolus dilution during transport through the vessel tree. The equilibrium contrast agent concentration, *b*, depends on the patient’s blood volume. Neglecting the filtration of contrast agent in the kidney, and contrast agent leakage, the upper bound for *b* is the administered amount of contrast agent divided by the total blood volume. However, *b*, can become lower in regions of contrast agent leakage and is dependent on the severity of the leakage and the size of the affected region in the brain. Here, we choose values for *a, b*, and *t_p_* within large enough bounds to ensure physically meaningful inflow profiles. The parameters *k_B_* and *k_T_* are dimensionless scaling factors for the effect of meso-scale 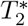-shortening due to the magnetic susceptibility contrast at the interface of the vascular and the extra-vascular, extra-cellular compartment and the interface of the extra-vascular, extra-cellular and the cell compartment, respectively. Because these values depend on the tissue architecture, *k_B_* and *k_T_* can also mitigate errors in the NMR signal prediction caused by patient-specific variations in vessel geometry. The pre-contrast relaxation times *T*_1,pre_ and 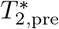 vary from voxel to voxel. From Eq. (17), it is clear that 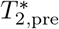, cancels when *S*(*t*) is normalized. Therefore, the value of 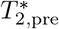 is not critical for the present study. The authors of [61] measured *T*_1,pre_ in patients with relapsing-remitting MS for several lesion types. They reported values between 1.9 s for black holes, and 0.8 s for NAWM, at 3 T. The filtration coefficient *L_p_* characterizes the fluid mass exchange between the vascular and the extra-vascular compartment. The authors of [45] suggest *L_p_* = 2.7 · 10^−12^ m Pa^−1^ s^−1^ for normal subcutaneous and *L_p_* = 2.1 · 10^−11^ mPa^−1^ s^−1^ for tumor tissue. While in normal brain tissue the contrast agent stays in the blood stream, it leaves the vascular compartment over the vessel wall in regions where the BBB is impaired. Therefore, the filtration coefficient Lp is likely to be elevated in such tissue, due to opened tight junctions. The diffusive capillary wall conductivity, *D_ω_*, characterizes the diffusive transport of contrast agent between the vascular and the extra-vascular compartment. It depends, i.a., on the molecular diffusion coefficient of the contrast agent, the wall thickness, porosity and the tortuosity of the transmural pathway.

### Parameter estimation

In a preliminary model investigation, we use the parameter estimation toolbox PEST [62] to find the parameter set that minimizes the sum of squared differences, 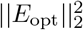, between the simulated signal-time curve and the MRI data. For the parameter estimation, we employ the truncated singular value decomposition algorithm, available in PEST. The estimated parameter values for the best fit against the curves N and L, cf. Fig. 1, as well as the corresponding ║*E*_opt_║_2_, are given in Table 1.

**Table 1.**
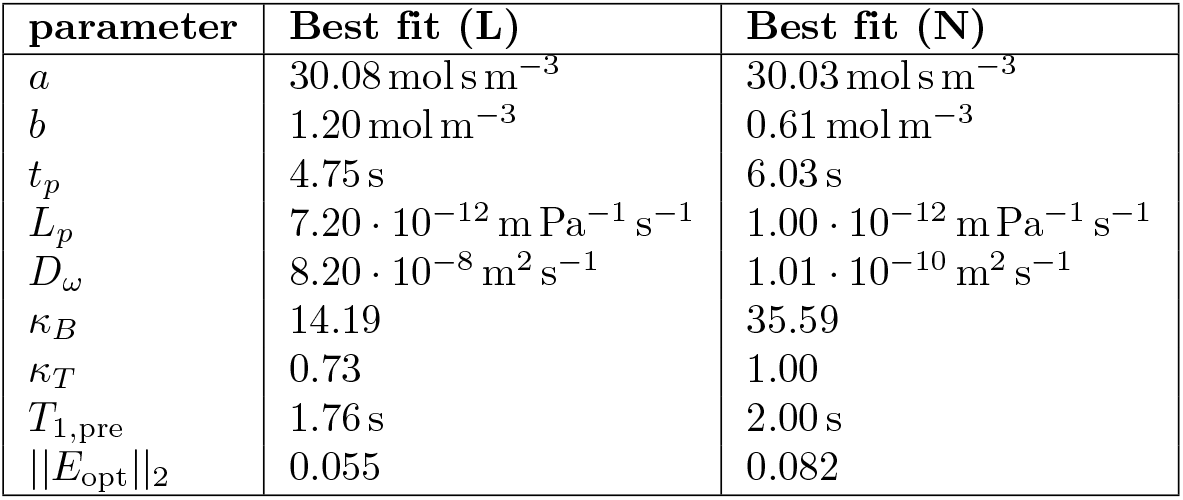
Parameter values obtained by a global optimization algorithm for the best fit of model and MRI data, minimizing ║*E*_opt_║_2_. The second column shows the parameters for the lesion sample (L), the last column the parameters for the NAWM sample (N).

| parameter | Best fit (L) | Best fit (N) |
| --- | --- | --- |
| $a$ | $30.08 \text{ mol s m}^{-3}$ | $30.03 \text{ mol s m}^{-3}$ |
| $b$ | $1.20 \text{ mol m}^{-3}$ | $0.61 \text{ mol m}^{-3}$ |
| $t_p$ | 4.75 s | 6.03 s |
| $L_p$ | $7.20 \cdot 10^{-12} \text{ m Pa}^{-1} \text{ s}^{-1}$ | $1.00 \cdot 10^{-12} \text{ m Pa}^{-1} \text{ s}^{-1}$ |
| $D_\omega$ | $8.20 \cdot 10^{-8} \text{ m}^2 \text{ s}^{-1}$ | $1.01 \cdot 10^{-10} \text{ m}^2 \text{ s}^{-1}$ |
| $\kappa_B$ | 14.19 | 35.59 |
| $\kappa_T$ | 0.73 | 1.00 |
| $T_{1,\text{pre}}$ | 1.76 s | 2.00 s |
| $\ E_{\text{opt}}\ _2$ | 0.055 | 0.082 |

A comparison of the simulated and measured NMR signals, Fig. 4, indicates that the model can reproduce the measured curves well. Table 1 shows that the diffusive wall conductivity, *D_ω_*, is estimated to be low for the NAWM sample (N), and high for the lesion sample (L), with a difference of three orders of magnitude, while the other parameters are within the same order of magnitude. To better understand the influence of the diffusive wall conductivity on the computed NMR signal, we compute the mass of contrast agent in the extra-vascular space

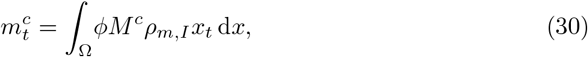

at the end of the simulation, *t*_end_ = 112 s. Additionally, we compute the total mass of contrast agent going into the domain over the entire time of the simulation,

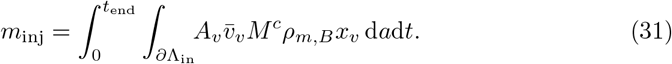

**Fig 4.**
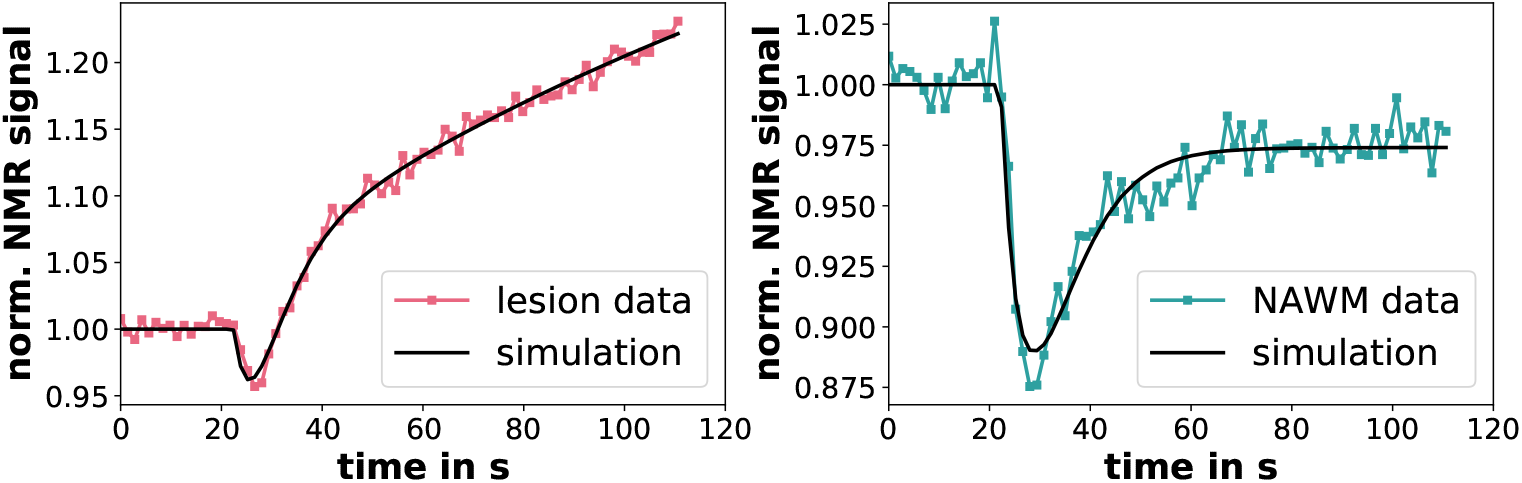
Simulated normalized NMR signals compared with MRI data (see Fig. 1), using the best fit parameter estimates given in Table 1. Left – the result for the lesion sample (L), right – the result for the NAWM sample (N).

The results are shown in Fig. 5 for different wall diffusivities. The other parameters were chosen as in Table 1, sample L. It can be seen that for *D_ω_* < 1.0 · 10^−9^ m s^−1^, there is almost no leakage into the extra-vascular space, i.e. the BBB is intact. For *D_ω_* > 3.0 · 10^−6^ ms^−1^, the leakage of contrast agent into the extra-vascular space has reached a plateau and does not increase further with *D_ω_*. For such high wall diffusivities, the contrast agent mole fractions in vascular and extra-vascular space reach an equilibrium. This situation would lead to a flat NMR signal (as seen, for instance, in the uppermost curve in Fig. 6 for *D_ω_*), which is not observed in any of the clinical data. Therefore, such high values of Dω are unlikely to be physiologically sensible. For the values of *D_ω_* in Table 1, this means that there is little to no contrast agent leakage for sample N, while there is significant leakage for sample L. This is in accordance with the present understanding of the pathology, which assumes leaky vessel walls in MS lesions.

**Fig 5.**
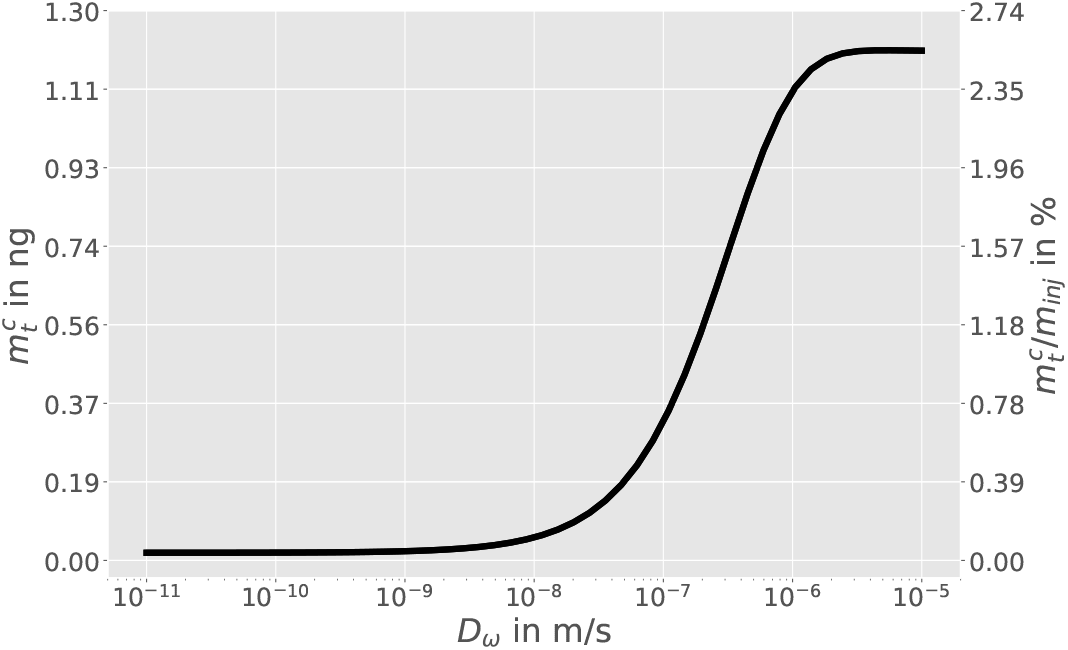
The mass of contrast agent in the extra-vascular space at *t*_end_ = 112 s for different wall diffusivities. The left axis shows the contrast agent mass in the extra-vascular space, 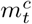. The right axis shows the ratio of 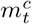 to the total injected contrast agent mass *m_inj_* in percent.

However, the problem of finding best fit parameters is typically ill-conditioned, or even ill-posed as the solution may be non-unique, such that the employed parameter estimation method may not be reliably applied. Therefore, we discuss other methods to further analyze the model parameters in the subsequent sections.

### Parameter sensitivity

For a better understanding of the influence of the patient-specific parameters on the signal-time curve, as well as the sensitivity of the model output to the model input parameters, we perform a simple sensitivity analysis, where parameters are individually varied, while all other parameters are kept constant at the values listed in Table 1. The results of this study are shown in Fig. 6 for sample L, and in Fig. 7 for sample N. It can be seen that the parameter sensitivity is different for L and N (which correspond to different locations in the parameter space). Such behavior characterizes non-linear model response.

**Fig 6.**
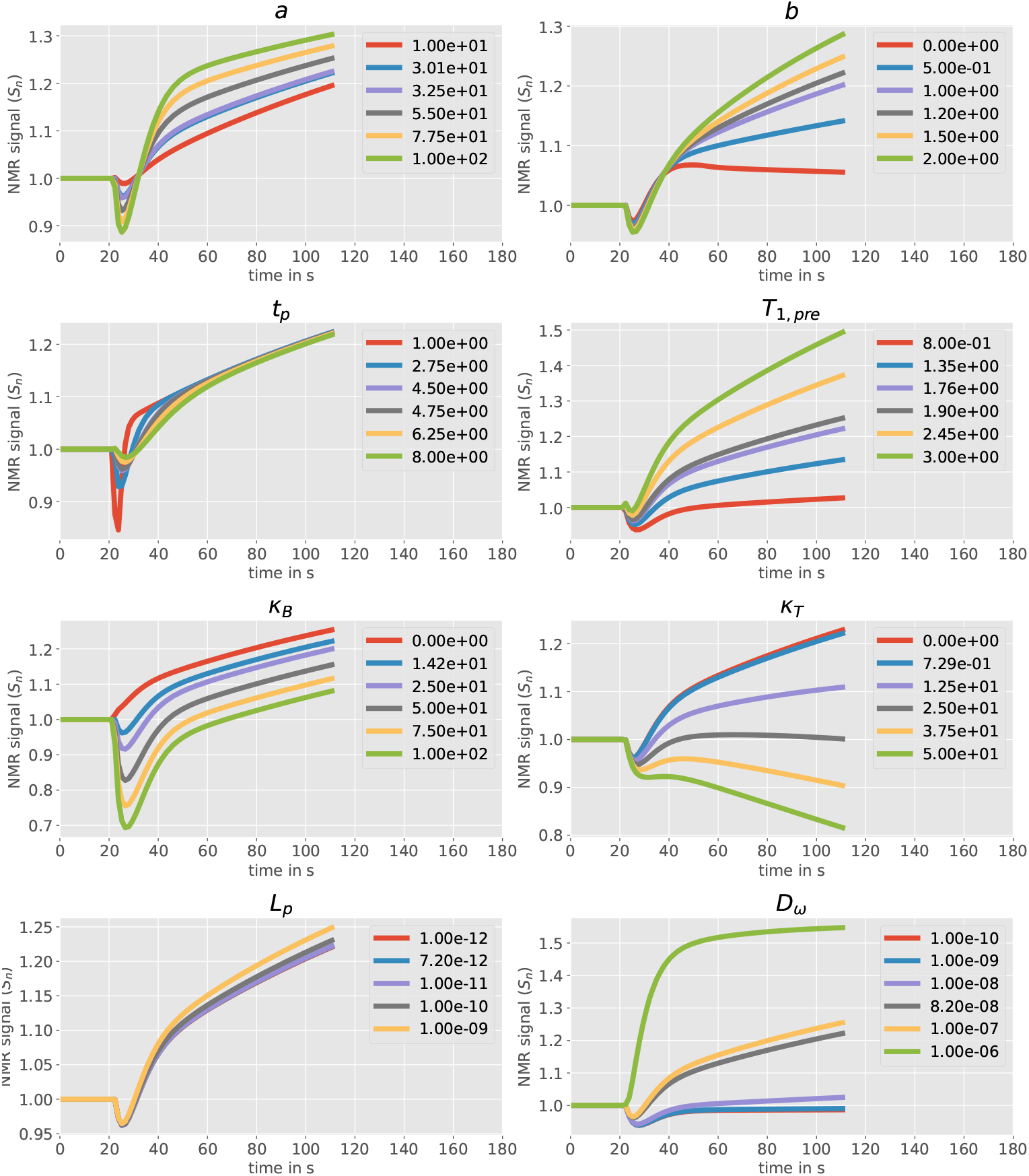
Influence of different flow, transport, and NMR parameters on the signal–time curve. The parameters are individually varied, while the other parameters are chosen as in Table 1 (sample L).

**Fig 7.**
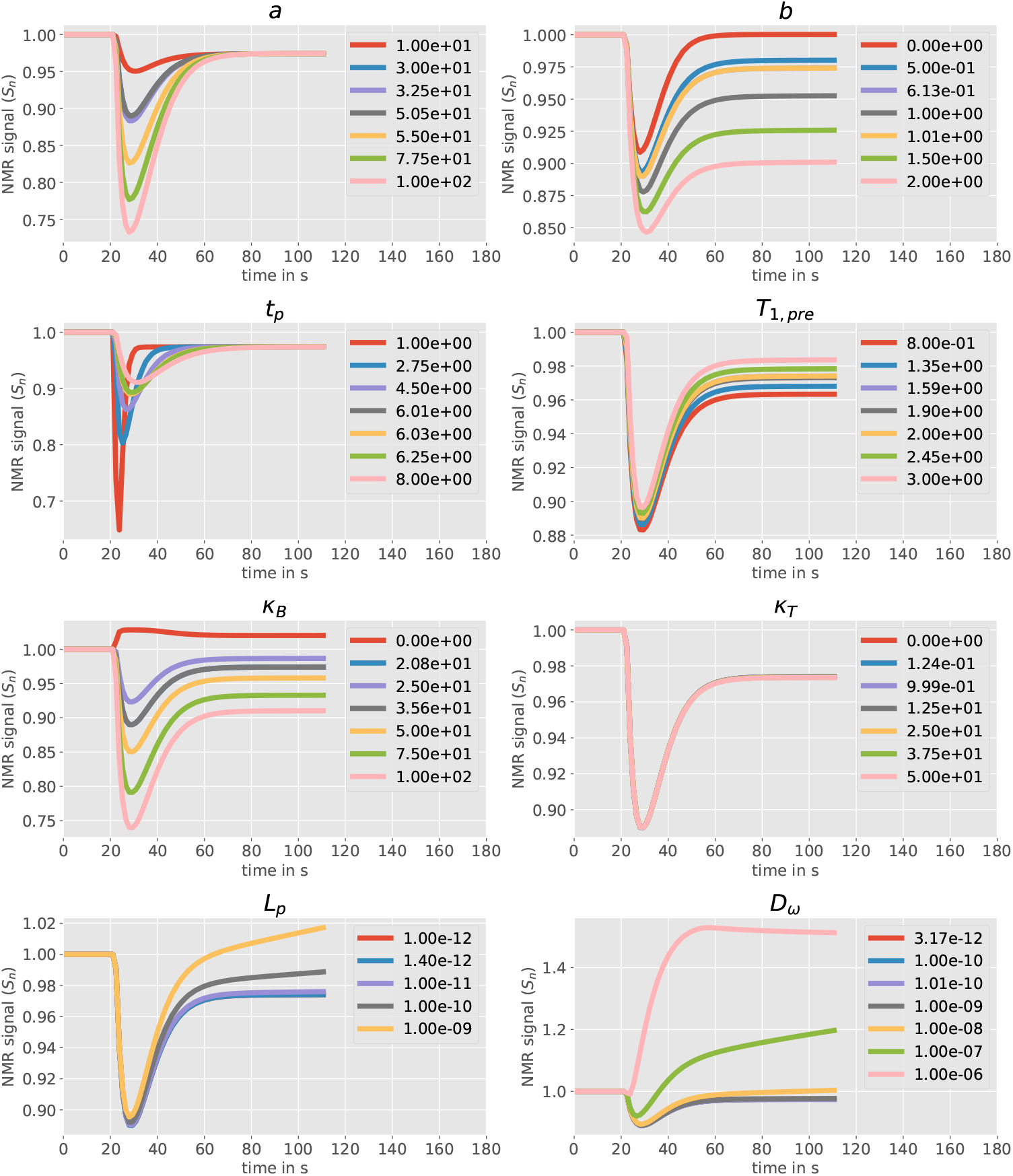
Influence of different flow, transport, and NMR parameters on the signal–time curve. The parameters are individually varied, while the other parameters are chosen as in Table 1 (sample N).

#### Capillary input function

The shape parameters *a* and *t_p_* of the capillary input function have a strong influence on the first pass dip of the NMR signal. The influence is directly related to the 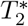-shortening caused by the contrast agent in the blood vessels. Comparing the respective curves in Fig. 6 and 7, shows that contrast agent leakage dampens the influence of a and *t_p_*. The difference in concentration between the vascular and extra-vascular space decreases in the presence of leakage, attenuating the 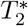-shortening meso-scale effects. For sample L, *a* also influences the signal in later times in the presence of leakage. A higher a indicates a larger contrast agent bolus, which will also result in a higher amount of leakage leading to a signal increase at later times, due to the *T*_1_-shortening effect of the contrast agent in the extra-vascular space. In the absence of leakage (sample N), the late signal is only affected significantly by the equilibrium concentration *b*. For sample L, *b* has a significant influence on the late signal slope. In that case, the signal slope is directly related to the leakage rate. With a higher *b*, the gradient of the contrast agent concentration over the vessel wall is higher, leading to a higher leakage rate. For *b* = 0, the slope is negative, indicating that leaked contrast agent flows back into the vascular compartment.

#### NMR parameters

The NMR parameters, *k_B_, k_T_, T*_1,pre_, have an equally strong but different effect on the NMR signal. The scaling parameter *κ_B_* for the meso-scale 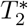-effects from the vascular wall, affects the signal strength almost linearly throughout the entire simulation. For *κ_B_* = 0, i.e. if meso-scale effects on 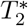-relaxation are neglected, the early time signal enhancement due to *T*_1_-shortening becomes even stronger than the signal decrease due to 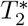-shortening, as clearly seen in Fig. 7. This illustrates that it is essential for the NMR signal model to include meso-scale effects. The scaling parameter *κ_T_* for the meso-scale 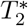-effects from the cell walls, only influences the signal in the presence of leakage (sample L). This is evident, since the difference between the contrast agent concentration in the cells and the extra-vascular, extra-cellular compartment is zero, in the absence of leakage. Fig. 6 shows that signal decrease due to 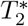-shortening in the extra-vascular compartment exceeds signal enhancement due to *T*_1_-shortening, if *κ_T_* is chosen too large. Because this is not seen in any of the clinical data, *κ_T_* is likely to be small (*κ_T_* < 10). The pre-contrast longitudinal relaxation time, *T*_1,pre_, shows a direct influence on the signal-enhancing effect of *T*_1_-shortening. If *T*_1_ is already elevated before the administration of contrast agent, the *T*_1_-shortening has a strong signal-enhancing effect. If *T*_1,pre_ is closer to *T*_1_ values measured for NAWM [61], the signal-enhancing effects are significantly weaker. Fig. 6 suggests that signal enhancement is small if *T*_1,pre_ is not elevated, even in the presence of leakage.

#### Leakage coefficients

The leakage coefficients for advective and diffusive transmural transport, *L_p_* and *D_ω_*, show a very similar qualitative influence on the NMR signal. However, the sensitivity of the NMR signal with respect to changes in *L_p_* is significantly lower than the sensitivity with respect to changes in *D_ω_*. This suggests that the main mechanism for transmural contrast agent leakage is of diffusive nature. Furthermore, note that changing *D_ω_*, while keeping the other parameters constant, can change the signal-time curve from the shape of sample N to the shape of sample L, and vice versa. This further emphasizes that diffusive wall conductivity plays a dominant role in characterizing curve shapes.

### Bayesian parameter inference

To complete our critical assessment of the proposed model, we ask and attempt to answer the question: *What can we learn about the model parameters, given the MRI data?* Bayesian parameter inference is a method to estimate unknown parameters of a model, given some prior knowledge about the parameters, and observations, while quantifying the uncertainty that is inherent to such a parameter estimation. Let *θ* denote the parameters of the model 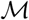, and *X* the vector of observed values. Bayes’ theorem, applied to the problem of parameter inference, states that

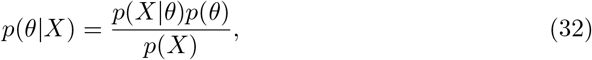

where *p*(*θ*|*X*) is the *posterior distribution*, i.e. the probability of *θ* given the observation data *X*. *p*(*X*|*θ*) is the *likelihood function*, i.e. the probability of the *X* being from the same population as the model prediction, given *θ*. *p*(*θ*) is the *prior distribution* reflecting prior knowledge about the parameters *θ*, before knowing the observations. *p*(*X*) is the *marginal likelihood*, a normalization constant, not depending on *θ*. Now, let 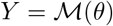 be the model prediction given the the parameters *θ*. We assume that we can write

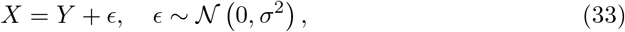

where *ϵ* is the combination of measurement error and unbiased model error and *σ* its standard deviation. The likelihood that any model answer, *Y*, comes from the same population as the measurement, *X*, is a Gaussian likelihood

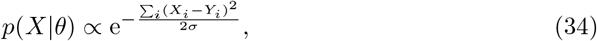

if the errors of all observations are assumed to be uncorrelated. The standard deviation, *σ*, has to be estimated for the given MRI data and the proposed model. We assume that our model represents the underlying physical processes accurately, so that model and discretization error are negligible compared to the MRI data measurement error. The measurement error is estimated from the MRI data obtained before the contrast agent bolus reaches the tissue sample, where the measurement is assumed to fluctuate around a constant baseline signal. To this end, we take 100 random signal samples from the brain slice shown in Fig. 1, normalize the signal to the mean of the first 10 sample data points, and compute the standard deviation of all such baseline data points across all samples, yielding *σ* = 0.009.

Markov chain Monte Carlo (*MCMC*) methods are methods to sample from the posterior distribution *p*(*θ*|*X*) without the need to compute marginal likelihood, which is generally expensive. MCMC draws samples on a random walk through the parameter space, creating a representative set of samples from the posterior distribution, after a sufficient number of iterations. These samples form a Markov chain such that the parameters with which the sample is generated in one step only depend on the parameters in the previous step. Herein, we use the ensemble sampler proposed in [63], which is implemented in the Python module *emcee* [64]. Its algorithm features an ensemble of interdependent Markov chains (so called walkers), enabling multiple parallel forward model runs within one step. The algorithm is briefly described in S2 Appendix. We refer to the literature [63, 64] for a comprehensive discussion.

In the following, Bayesian parameter inference is used to compute the probability distribution of the patient-specific model parameters, under physical parameter constraints, given a signal-time curve from a voxel of a perfusion MRI sequence. To this end, we choose the prior distributions of the parameters to be uniform distributions within the bounds given in Table 2. The ensemble sampler is configured with *k* = 100 walkers. The parameter vector is *θ* = [*a, b, t_p_*, log_10_ *D_ω_*, *T*_1,pre_, *k_B_, k_T_*]^*T*^, so that *N* = 7. The parameter *L_p_* remains fixed to reduce the dimension of the parameter space. Its influence on the NMR signal has been shown in the previous section to be significantly weaker than the influence of *D_ω_* (see Fig. 6).

**Table 2.**
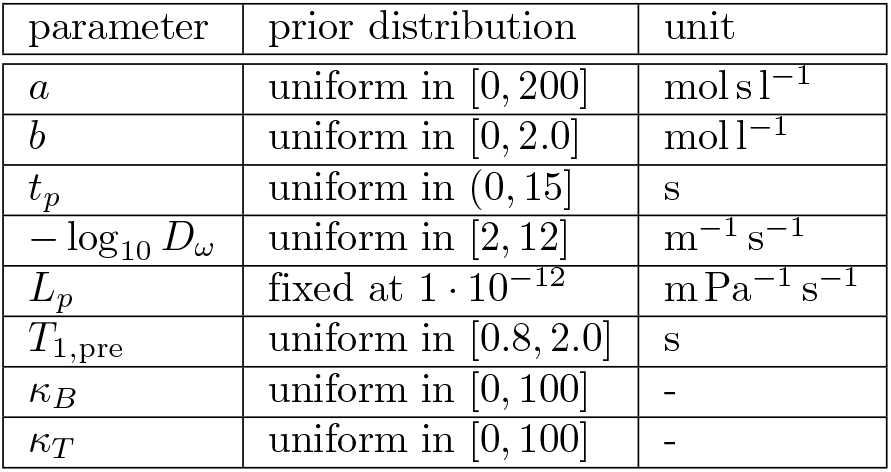
Prior distribution for parameters inferred by a Markov chain Monte Carlo method.

The sampler convergence is estimated using the integrated auto-correlation time, *τ_f_* [63]

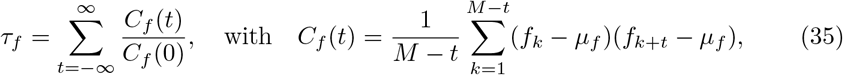

where 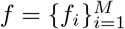 is a finite chain of length *M*, e.g. the value of parameter *a* for each sample in the Markov chain, and *μ_f_* its arithmetic mean. We use an estimate of the integrated auto-correlation, *τ_f,e_*, using the Python module *acor* [65, 66]. We compute this estimate for the chain of each parameter, *θ_i_*, and use the minimum and maximum values, *τ*_max_ = max_0≤*i*<*N*_ *τ_θ_i_,e_*, *τ*_min_ = min_0≤*i*<*N*_ *τ_θ_i_,e_*. The sampler is run until the sample size, *j* > 100 · *τ*_max_, and the change in the auto-correlation time estimate from sample *j* − *τ*_max_ to sample *j* is less than 1 %. The resulting histograms for each parameter and their covariance with respect to the other parameters is visualized in Fig. 8 for sample L and Fig. 9 for sample N (cf. Fig. 1). To eliminate artifacts from the burn-in phase of the MCMC algorithm, the first 10 · *τ*_max_ samples are discarded. To have only independent samples, every *τ*_min_ sample of the remaining samples is chosen [63], while the others are discarded. The solid black lines in Figs. 8 and 9 show the parameter values of Table 1 that were obtained previously with PEST.

**Fig 8.**
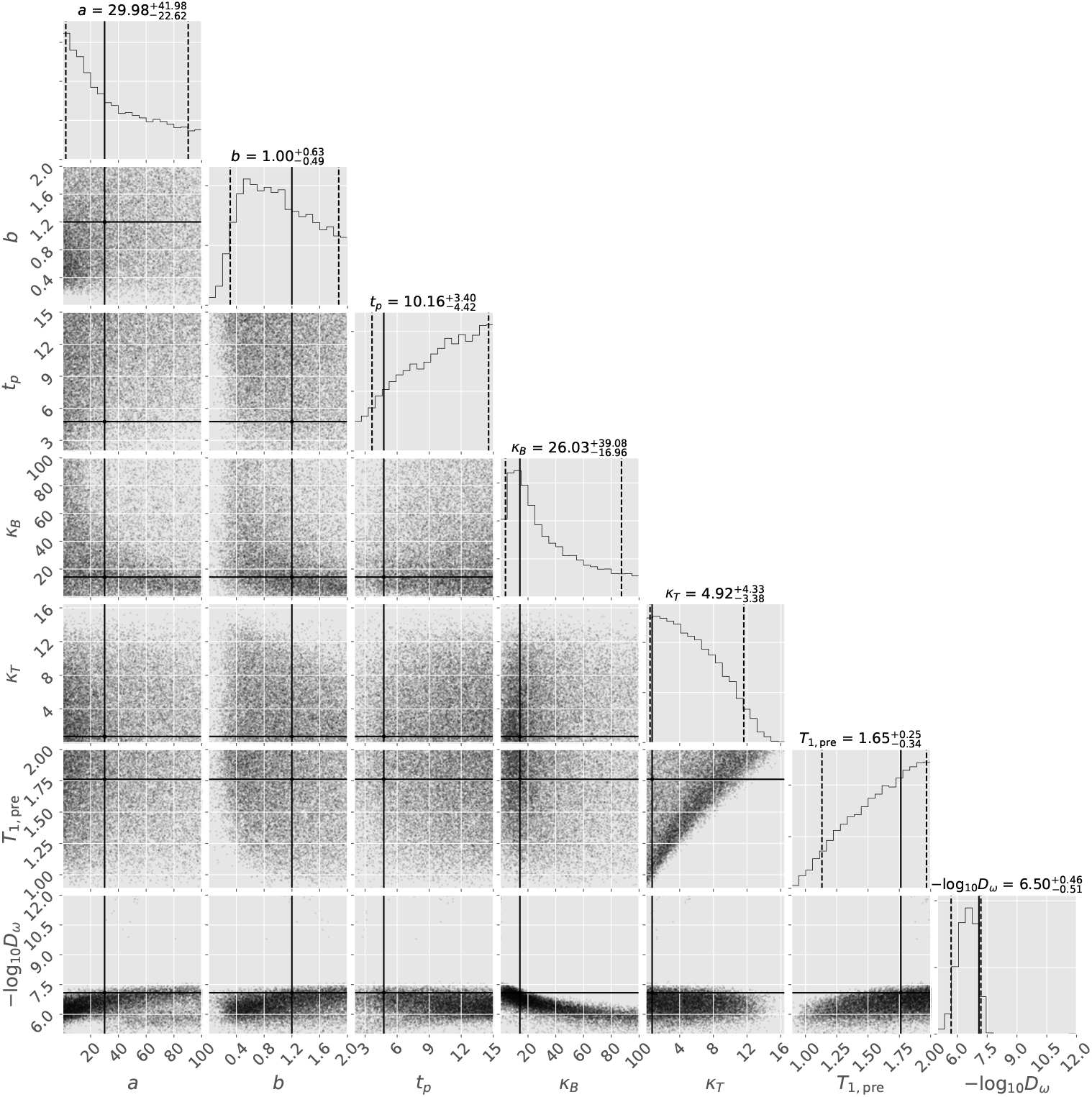
Histograms of model parameter distributions after learning from MR voxel data from a lesion (see Fig. 1, sample L). The histograms on the diagonal are the histograms for single parameters, the scatter plot in the matrix shows the covariance between the respective row and column parameters (plot generated with [67]). The histogram titles show median, 5th, and 95th percentile (also visualized as dashed lines). The horizontal and vertical solid black lines show the parameter values for sample L of Table 1.

**Fig 9.**
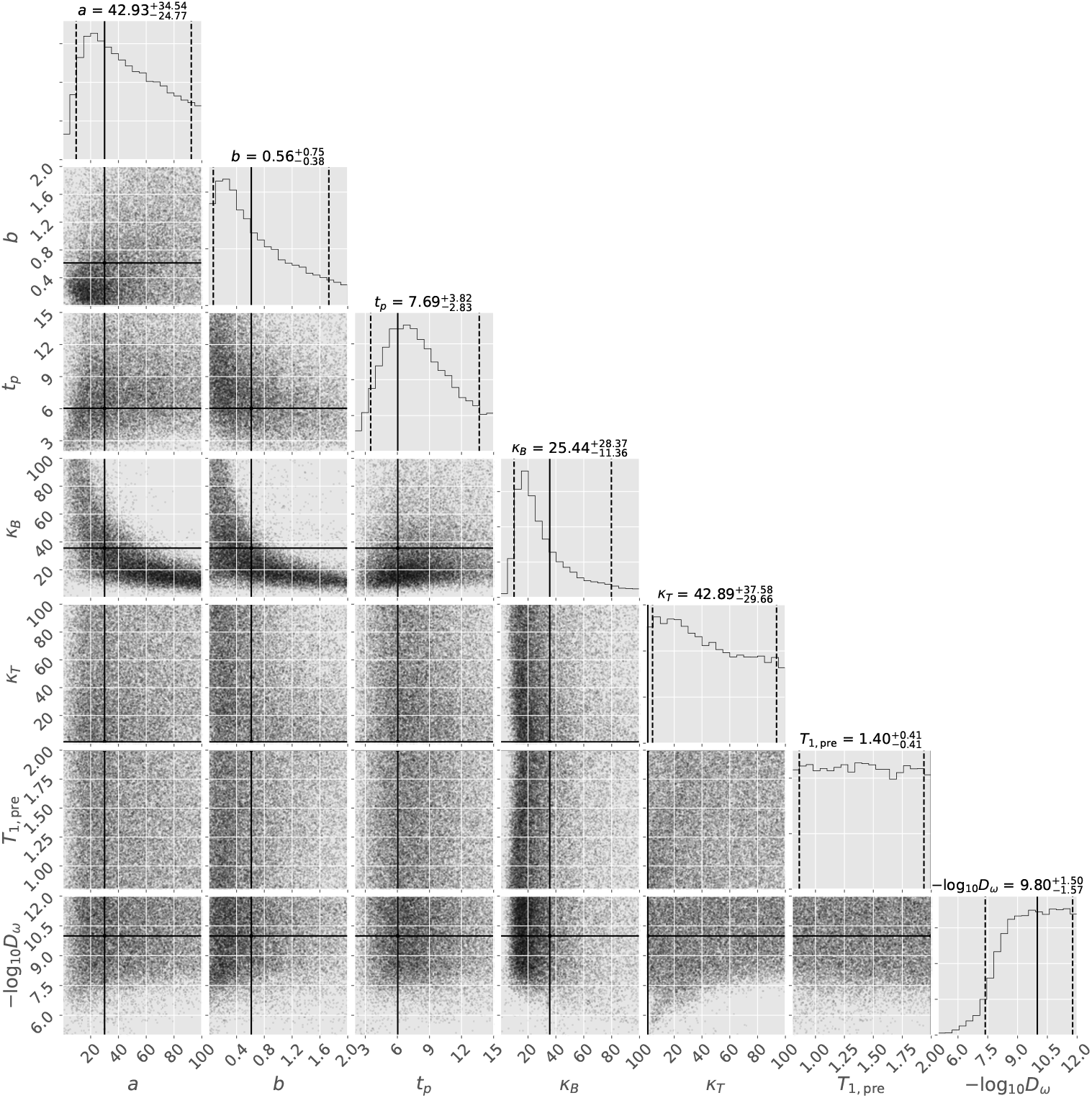
Histograms of model parameter distributions after learning from MR voxel data from NAWM (see Fig. 1, sample N). The histograms on the diagonal are the histograms for single parameters, the scatter plot in the matrix shows the covariance between the respective row and column parameters (plot generated with [67]). The histogram titles show median, 5th, and 95th percentile (also visualized as dashed lines). The horizontal and vertical solid black lines show the parameter values for sample N of Table 1.

To interpret the results, we recall the original question: *What can we learn about tht model parameters, given the MRI data?*

If the posterior distribution of a parameter is close to uniform, i.e. close to the prior distribution (see Table 2), the data did not provide any additional information about this parameter. This is the case for *κ_T_* and *T*_1,pre_ in Fig. 9, which is consistent with the observation in Fig. 7 that the sensitivity of the NMR curve with respect to changes in *k_T_* or *T*_1,pre_ is low.

In contrast, if the posterior distribution differs significantly from the prior distribution, the data provides significant information on this parameter. This is the case for the parameters *D_ω_* and *κ_T_* in Fig. 8, and *D_ω_* and *κ_B_* in Fig. 9. Again, this is consistent with the observation in Figs. 6 and 7 that the sensitivity of the NMR curve with respect to those parameters is high, such that only a small range of values for those parameters is likely to match the model results with the clinical MRI data.

Most interestingly, the distribution of *D_ω_* in Fig. 8 differs significantly from the distribution of *D_ω_* in Fig. 9. Both distributions are shown as histograms in Fig. 10. For sample N, the inferred diffusive wall conductivity is very likely to be below *D_ω_* = 1 · 10^−9^ ms^−1^, while all smaller values are equally likely. The magnitude of the value suggests that diffusive transport of contrast agent across the capillary wall is negligible (see Fig. 5). For sample L, the inferred distribution diffusive wall conductivity, has a distinct peak around *D_ω_* = 4 · 10^−7^ ms^−1^. Furthermore, it shows that values below *D_ω_* = 3 · 10^−8^ms^−1^ are very unlikely, suggesting significant transmural contrast agent leakage. It can be concluded that the two samples can be distinguished just on the basis of *D_ω_*, without looking at the estimates for the other parameters.

**Fig 10.**
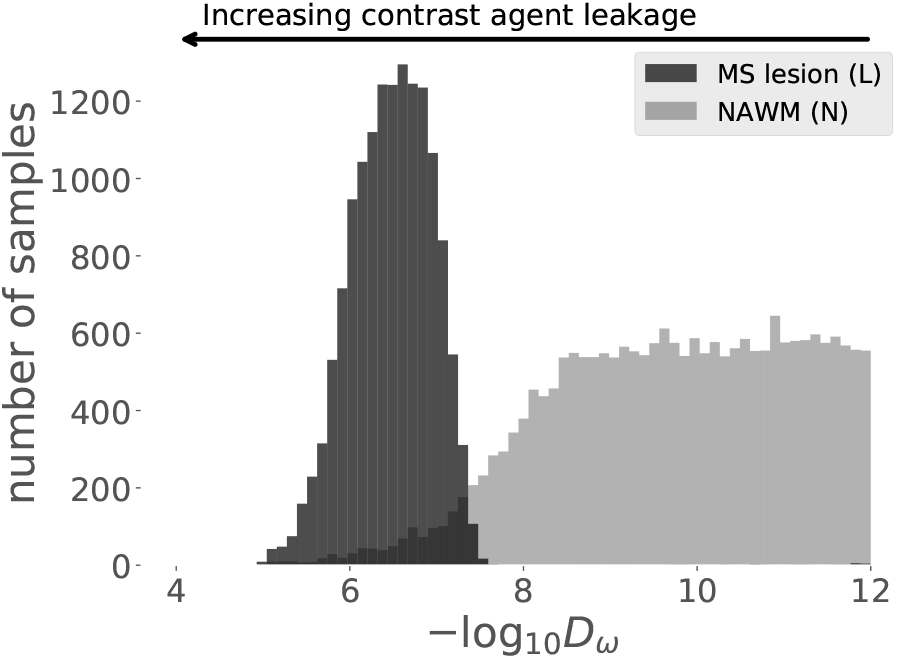
Histograms for Bayesian parameter inference, when learning from NAWM data or contrast-enhancing lesion data. A low diffusion coefficient is most likely for the NAWM data, while a high diffusion coefficient is most likely for the contrast-enhancing lesion data.

The uncertainty in *D_ω_* reflects the fact that all other parameters are uncertain as well. Consequently, the estimate of *D_ω_* may be improved with additional information about other parameters. Such information might be, for instance, a direct measurement of *T*_1,pre_, estimations of the AIF, or data from other MR sequences of the same patient. Furthermore, knowledge that a parameter is expected to be similar in a certain region of the brain, could enable learning from other voxel data of the same sequence. In the Bayesian framework, such information can be included incrementally, where the posterior distributions of the previous Bayesian update are the prior distributions of the next Bayesian update.

## Model limitations and outlook

The current model relies on a single exemplary vessel geometry. Today, patient-specific sub-voxel vessel geometries cannot be routinely measured. Hence, the influence of different vessel geometries on the presented results has to be investigated.

Furthermore, the used model of the inflow curve (Eq. (14)) neglects re-circulation in the form of a second or third pass of the contrast agent. In particular, the effect of the second pass of the bolus cannot be captured and might lead to more uncertainty in the estimation of other model parameters. In a future step, the inflow curve model can be improved to include re-circulation and to be derived from AIF measurements.

The presented model considers processes in a sub-voxel tissue sample that is surrounded by tissue with the same properties. However, contrast-enhancing lesions in the brain typically span over several MRI voxels, see Fig. 1. Furthermore, patterns such as ring-like shapes have been observed for MS [68], suggesting processes on a larger scale, or possible inter-voxel dependencies. Such effects can not be included in the model in its current state, since simulation of several voxels are prohibitively expensive due to the large number of blood vessels.

The applicability of the presented model has yet to be confirmed in a clinical environment. This would be of special relevance for monitoring of pharmacologic effects and drug efficacy, e.g. in drugs that are targeted against immune cell trafficking. It is to be analyzed how reliable the method predicts diffusive capillary wall conductivities over a wider range of patient-specific data.

A current drawback of the method is the computational time required to infer diffusive capillary wall conductivities and contrast agent leakage. However, the computational cost can most likely be improved by applying model reduction techniques and machine learning algorithms. Likewise, homogenization techniques can be used for model reduction [69, 70]. However, such techniques are difficult to apply, due to the hierarchical structure of the micro-circulation. For all approaches, the presented model can be used as theoretical basis and as validation tool.

## Summary and conclusion

We presented a mixed-dimension fluid-mechanical model for contrast agent brain tissue perfusion on the sub-voxel scale. The blood vessels are considered as a network of cylindrical tubes. The extra-vascular compartment is modeled as a porous medium. The presented discretization results in a coupled system of partial differential equations of three-dimensional and one-dimensional equations. The fluid-mechanical model can describe the three-dimensional evolution of the contrast agent concentration on the sub-voxel scale. We further proposed an NMR signal model, describing the influence of the contrast agent on the NMR voxel signal, including meso-scale effects. A convergence study suggests that the combined model is consistent and converges to a unique solution on grid and time step refinement. Using parameter estimation, it was shown that the model can describe two characteristic NMR signal curves from clinical data obtained by DSC-MRI for a patient with MS lesions, and that the estimated model parameters provide a meaningful physical interpretation. Bayesian parameter inference, with the given model and clinical DSC-MRI data, showed that the two given NMR signal curves can be distinguished and characterized, only on the basis of the estimated diffusive capillary wall conductivity distributions. The study suggests that the NMR signal curve, given the model, is informative about some patient-specific model parameters, such as the diffusive capillary wall conductivity, and less informative about others, such as the tissue’s *T*_1_ relaxation time before contrast agent administration. Furthermore, the uncertainty of the diffusive capillary wall conductivity predictions could be quantified in the Bayesian framework. In summary, the presented model constitutes a useful tool to study contrast agent perfusion on a sub-voxel scale, and may lead to an improved understanding of the sub-voxel processes beyond the scope of this paper.

## Supporting information

### S1 Appendix. Numerical convergence study in space and time

As there is no analytical solution to the problem, we use the solution computed with a very fine resolution in space and time as a reference for a grid convergence study. We compute errors with respect to the fine scale solution by mapping the coarse scale solution to the finest grid. Let 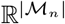 be the solution space on grid 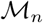, where 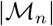 is the number of cells on grid level *n*, and 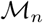 is a set of hexahedra, *K_i_*, such that 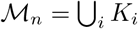 is a discrete representation of Ω. The levels are constructed by dividing each hexahedron on level *n* into 8 hexahedrons on level *n* + 1, such that 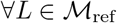, there exists exactly one 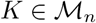 such that *L* ⊂ *K*, where 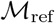 denotes the reference grid. The mapping from coarse to fine scale, 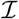, can be defined as

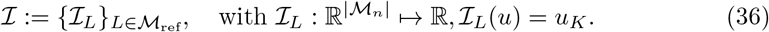

The relative error of the physical quantity *u* ∈ {*p_t_,x_t_*}, between reference and a solution on a coarser grid, *u*_ref_, *u*, respectively, is defined as

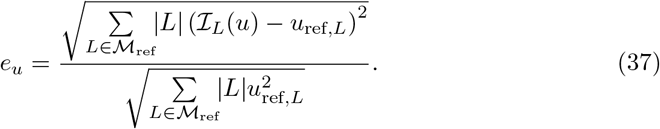

In time, we define the maximum relative error over all time steps *t^i^, i* ∈ {0, ⋯, *τ*}, where *τ* is the number of time steps, as

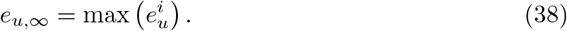

Finally, we measure the difference of the signal-time curve, *S*, to the reference curve, *S*_ref_, computed with the finest spatial and temporal discretization, in the following norm

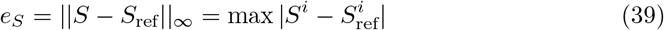

The convergence rates for a given error *e* are computed from one refinement level *n* to the next as

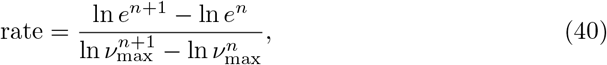

where *ν*_max_ is the respective maximum discretization length. In space, *ν*_max_ is defined as the maximum edge length of all elements, h_max_. When refining, the vessel domain grid is also refined by bisecting large elements until the maximum element length is smaller than *h*_max_. In time, *ν*_max_ is defined as the maximum time step size. The time step size, Δ*t*, is chosen to be small around the time where the contrast agent front reaches the domain, and increasingly larger as the process becomes slower, following the heuristic

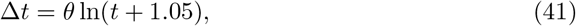

where *θ* > 0 is a factor controlling the time step size in the refinement study.

The reference solution is obtained with *h*_max_ = 1 μm and *θ* = 0.125. The parameters are chosen to be the optimal parameter set computed by an optimization algorithm described in the following section *Parameter estimation* (see Table 1), minimizing the signal difference to the MRI data from an MS lesion shown in Fig. 1 (in red).

Table 3 show the errors and convergence rates of the extra-vascular fluid pressure, *p_t_*, and the contrast agent mole fraction, *x_t_*. Fig. 11 shows the NMR signal curves and errors with respect to the reference solution when refining in space and time.

**Table 3.** Errors and convergence rates in space for the pressure, *p_t_*, and the mole fraction of the contrast agent, *x_t_*, in the extra-vascular domain.

| $h_{\max}$ | $e_{p_t}$ | rate | $e_{x_t, \infty}$ | rate |
| --- | --- | --- | --- | --- |
| 32 $\mu\text{m}$ | 0.000414833 | - | 0.028418 | - |
| 16 $\mu\text{m}$ | 0.000210479 | 0.978853 | 0.0152967 | 0.893589 |
| 8 $\mu\text{m}$ | 0.000107219 | 0.973115 | 0.00874756 | 0.806266 |
| 4 $\mu\text{m}$ | 5.31281e-05 | 1.01302 | 0.00471859 | 0.890526 |
| 2 $\mu\text{m}$ | 2.54764e-05 | 1.06032 | 0.00228896 | 1.04366 |

**Fig 11.**
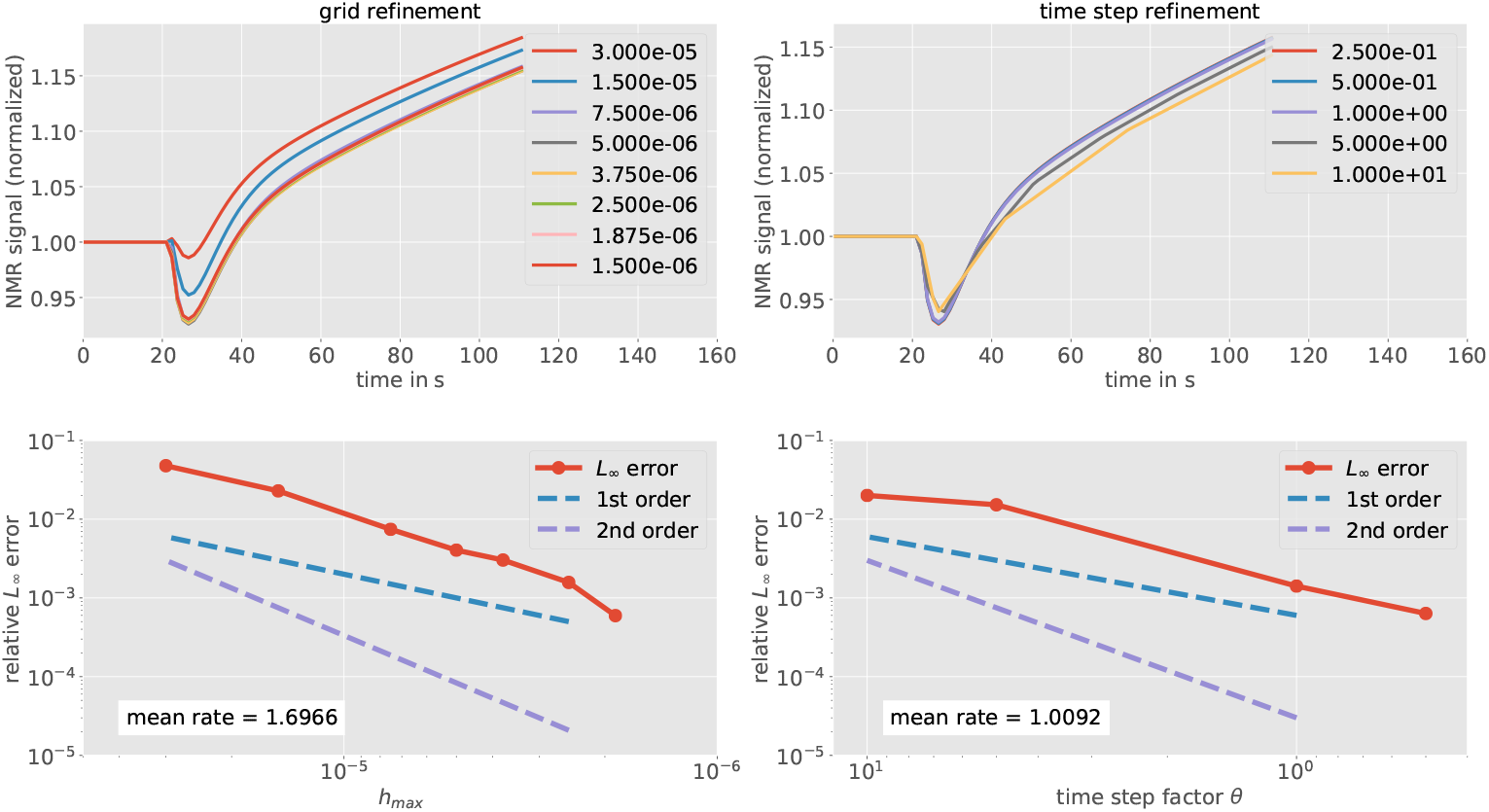
The NMR signal curves and errors to reference solutions when refining in time, while keeping the same fine resolution in space (left), and in space, while keeping the same fine resolution in time (right). The top left legend indicates the grid cell size in m, the top right legend indicates the time step factor *θ* from Eq. (41). The bottom left error plot shows convergence with a mean rate of 1.6966 with grid refinement. The bottom right error plot shows convergence with a mean rate of 1.0092 with time step refinement.

It can be seen that all quantities converge to the reference solution. We obtain convergence rates close to 1 for the pressure and the mole fraction of the contrast agent. The signal curve converges with first order in time and a slightly higher order in space. The higher convergence may be explained by the computation of the signal involving the integration of the concentration over the entire domain. The relative error with respect to the reference solution, is smaller than 1 % for a moderate spatial and temporal refinement. In conclusion, we consider a spatial resolution of *h*_max_ = 8 μm, and a temporal resolution *θ* = 1 as sufficient for the subsequent analysis. We justify this with the assumption that the errors resulting from model parameter uncertainty, as well as the errors in the measurement data, are larger than the discretization error. This is also evident, when looking at the results of the parameter study and comparing the variability with that of the signal-time curves shown in Fig. 11 for different spatial and temporal discretizations. In order to verify that the discretization error is small also for other parameter configurations, we ran the above analysis for various parameter configurations and confirmed that the analysis looks similar for those other cases. The results are omitted for brevity.

### S2 Appendix. Brief description of the ensemble sampler

The employed ensemble sampler [63] considers two sets of *k*/2 random walkers, *S*_0_ = {*w_i_*}_*i*=1,…, *k*/2_, *S*_1_ = {*w_i_*}_*i*=*k*/2+1,…, *k*_, where the position of walker *w_i_* in step *n* is a position in the parameter space (a vector of parameters), denoted as 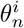. After each step the walkers in *S_m_* are moved such that

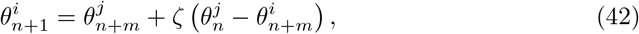

where *θ^j^* is a walker position randomly drawn from the positions of the other set of walkers, *S*_1−*m*_, and *ζ* is a random variable drawn from a proposal distribution *g*(*ζ*),

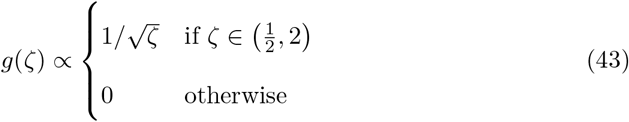

Note that this means moving the walkers in *S*_0_ first, then the walkers in *S*_1_. At each walker position, a sample is proposed. The sample is accepted with the probability [64]

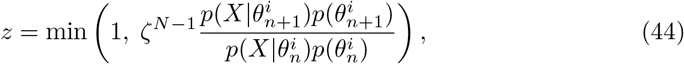

where *N* = dim(*θ*) is the dimension of the parameter space. If the sample is not accepted, the walker remains at the position 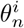, increasing the number of samples at this position by one. Each step requires a run of the forward model for every walker, which is computationally the most expensive part. Fortunately, advancing the walkers within a set of walkers can be done in parallel.

## Acknowledgments

The authors would like to thank V.G. Kiselev, University Medical Center Freiburg, for an interesting discussion about NMR physics and meso-scale effects, and M. Sinsbeck, University of Stuttgart, for helpful suggestions concerning Bayesian parameter inference.

